# Deriving Disease Modules from the Compressed Transcriptional Space Embedded in a Deep Auto-encoder

**DOI:** 10.1101/680983

**Authors:** Sanjiv K. Dwivedi, Andreas Tjärnberg, Jesper Tegnér, Mika Gustafsson

## Abstract

Disease modules in molecular interaction maps have been useful for characterizing diseases. Yet biological networks, commonly used to define such modules are incomplete and biased toward some well-studied disease genes. Here we ask whether disease-relevant modules of genes can be discovered without assuming the prior knowledge of a biological network. To this end we train a deep auto-encoder on a large transcriptional data-set. Our hypothesis is that such modules could be discovered in the deep representations within the auto-encoder when trained to capture the variance in the input-output map of the transcriptional profiles. Using a three-layer deep auto-encoder we find a statistically significant enrichment of GWAS relevant genes in the third layer, and to a successively lesser degree in the second and first layers respectively. In contrast, we found an opposite gradient where a modular protein-protein interaction signal was strongest in the first layer but then vanishing smoothly deeper in the network. We conclude that a data-driven discovery approach, without assuming a particular biological network, is sufficient to discover groups of disease-related genes.

## Introduction

Individualized treatment, i.e., precision medicine, is the holy grail of healthcare, since the majority of commonly used drugs work ineffectively for most patients, and the expected cost for developing a new drug currently exceeds $3 billion (Schork et al. 2015). This is probably due to the complexity of the many contributing disease factors. Moreover, individual markers for drug selection generally work poorly, and the choice of drugs in complex diseases is often based on trial and error strategies, causing suffering for patients and increasing costs for health care. To enable precision medicine, molecular networks derived from multiple omics such as transcriptomics, proteomics, genomics, and methylomics has been suggested to capture the complexity (Hood et al. 2015). Omics could potentially revolutionize medicine by analyzing disease as system perturbations rather than the malfunctioning of individual genes, thereby allowing for the application of systems medicine to complex diseases. Omics are routinely performed nowadays, and current data collections of different omics are rapidly growing and accessible. For example, UK Biobank offer researchers to analyze genotypes of ~500,000 indivduals (Bycroft et al. 2018), and several expression repositories contain data-sets exceeding 20,000 samples (e.g. Torrente et al. (2016), Lachmann et al. 2018). Systems medicine applications to date have often utilized the fact that disease genes are functionally related and their corresponding protein products are highly interconnected and co-localized within networks, thereby forming disease modules (Gustafsson et al. 2014; Menche et al., 2015). Several module-based studies have been performed on different diseases by us and others, forming the disease module paradigm (Barabási et al., 2011; Gustafsson et al. 2014; Menche et al., 2015, Hellberg et al. 2016). The modules generally contain many genes and a general principle for validation has been to use genomic concordance, i.e., the module derived from gene expression and protein interactions can be validated by enrichment of disease-associated SNPs from GWAS. The genomic concordance principle was also used in a DREAM challenge that compared different module-based approaches (Choobdar et al. 2019). Yet biological networks defining such modules are incomplete and biased toward some well-studied disease genes (Skinnider et al., 2018).

Deep artificial neural networks (DNNs) are revolutionizing areas such as computer vision, speech recognition, and natural language processing (LeCun et al. 2015), but only recently emerging to have an impact on systems and precision medicine (Topol 2019). For example, the performance of the top five error rates for the winners in the international image recognition challenge (ILSVRC) dropped from 20% in 2010 to 5% in 2015 upon the introduction of deep learning using pre-trained DNNs that were refined using transfer learning (Hinton et al. 2006). DNN architectures are hierarchically organized layers of non-linear machine learning algorithms (Deng et al. 2014). The layers in a deep learning architecture correspond to concepts or features in the learning domain, where higher-level concepts are defined from lower-level ones. Variational auto-encoders (AEs) represent a certain type of DNN that aims to mimic the input signal using a compressed representation, where principal component analysis represents the simplest form of a shallow linear AE. Given enough data, deep AEs have the advantage of being able both to create relevant features from raw data and identify highly complex non-linear relationships, such as the famous XOR switch, which is true if only one of its inputs is. Although *omics* repositories have increased in size lately, they are still several orders of magnitude smaller compared to image data sets used for ILSVRC. Therefore, effective DNNs should be based on as much omics data as possible, potentially using transfer learning from the prime largest repositories and possibly also incorporating functional hidden node representation using biological knowledge. The LINCS project defined and collected microarrays measuring only 1000 carefully selected landmark genes, which predicted 95% of the expression of all genes using DNN (Yifei et al. 2016). However, although interesting and useful for prediction purposes, those representations cannot be used for data integration or to interpret the DNNs. For that purpose, Tan et al. used denoising AEs derived from the transcriptomics of *Pseudomonas aeruginosa* and found a representation where each node coincided with known biological pathways (Tan et al. 2016). Chen et al. used cancer data and showed that starting from pathways represented as *a priori* defined hidden nodes, allowed the investigators to explain 88% of variance, which in turn produced an interpretable representation (Chen, et al., 2018). These results demonstrate that AEs can use pre-defined functional representations, and can learn such representations from input data, although systematic evaluation of how to balance between pre-defined features versus purely data-driven learning remains to be determined. To address this question, the interpretation of the representations within NNs is fundamental. The most commonly used tool for this purpose is the Google DeepDream (Mordvintsev et al., 2015). Briefly, the output effect is analyzed using the *light-up* of an activation of one hidden node, followed by a forward-propagation of the input to the output. This allows the user to interpret the net effect of a certain node and is referred to by us as *light-up*.

In this work, we investigated different AE architectures searching for a minimal representation that explained gene expression, which in our hands resulted in a 512-node wide and three-layered deep auto-encoder capturing ~95% of the variance in the data. Next, we derived a novel reverse supervised training-based approach based on *light-up* of the top of the transferred representations of the trained AE that defined the disease module. Using the third layer of the AE we identified disease modules for eight complex diseases and cancers which were all validated by highly significant enrichments of GWAS genes of the same disease. In order to understand the role of each of the hidden representations we tested whether they corresponded to genes that were functionally related and diseases associated genes. First, unsupervised analysis of the samples in the AE space showed that disease cluster in all layers, while cell types clustered only in the third layer. Then, we decoded the meaning of non-linear transformations that defined the compressed space of the auto-encoder. To this end we utilized the protein-protein interaction data in STRING (ver 9.1, Franceschini et al., 2013), as a first step to remove the essence of the knowledge of interactome in defining the phenotypic modules. Conversely, we found that genes within the same hidden AE node in the first layer were highly interconnected in the STRING network, which gradually vanished across the layers. In summary, we believe that our data-driven analysis using deep AE with a subsequent knowledge-based interpretation scheme, enables systems medicine to become sufficiently powerful to allow unbiased identification of complex novel gene-cell type interactions of relevance for realizing systems medicine.

## Results

### A deep auto-encoder with 512 hidden nodes accounts for 95% of the total variance of the data

Training neural networks requires substantial, well-controlled big data. In this study we therefore performed our analysis using the 27,887 quality-controlled and batch-effect corrected Affymetrix HG-U133Plus2 array compendium, thus encompassing data from multiple laboratories (Torrente et al., 2016). Furthermore, the data had previously been analyzed using cluster analysis and linear projection techniques such as principal component analysis (Torrente et al., 2016). The data set and the ensuing analysis therefore constitute a solid reference point based on which we are in a good position to ask whether successive non-linear transformations of the data would induce a biologically useful representation(s). Specifically, we investigate whether disease-relevant modules could be discovered by training an auto-encoder using this data set. The rationale is that by inducing identity mapping from input to output we can readily inspect the resulting deep representation from a disease module standpoint. To this end we partitioned the data into 20,000 training and 7,887 test samples. We trained AEs of different widths from 64 to 1024 hidden nodes, incremented stepwise in powers of two, and we contrasted two depths in our analysis, i.e. a *shallow* single-layered coded AE (SAE) and a *deep* triple layered AE (deepAE). We calculated the mean squared training and test error (measured using error = 1-R^2^, where R^2^ is computed globally over all genes using a global data variance (Fig. 1A) and locally for each gene individually using gene-wise variances (Fig. 1B-C)). Comparing the reconstruction errors of the different auto-encoders we found, not surprisingly, that the SAE performed poorly (>15% error) whenever we used less than 1024 hidden nodes, whereas increasing the number of nodes to 1024, reduced the error three-fold to ~5%. In contrast, the deepAE performed well already for 64 hidden nodes (11% error), which subsequently decreased following a power law up to 512 hidden nodes, best described by R^2^ = 0.89*2^0.028(x-64)^, where x is the number of hidden nodes. Next, we analyzed the gene-wise R^2^ performances of the R^2^ distributions (Fig. 1B-C) which showed that the median gene error was also low (R^2^ > 0.86) already for the 512 deepAEs. In summary, we found that the deepAE with 512 hidden nodes performed comparably to the SAE with 1024 nodes. Since the purpose of our study was to discover biologically meaningful disease module we proceeded and analyzed the 512 deepAEs in the remaining part of the paper as this architecture provided an effective compression of the data.

**Fig. 1.**
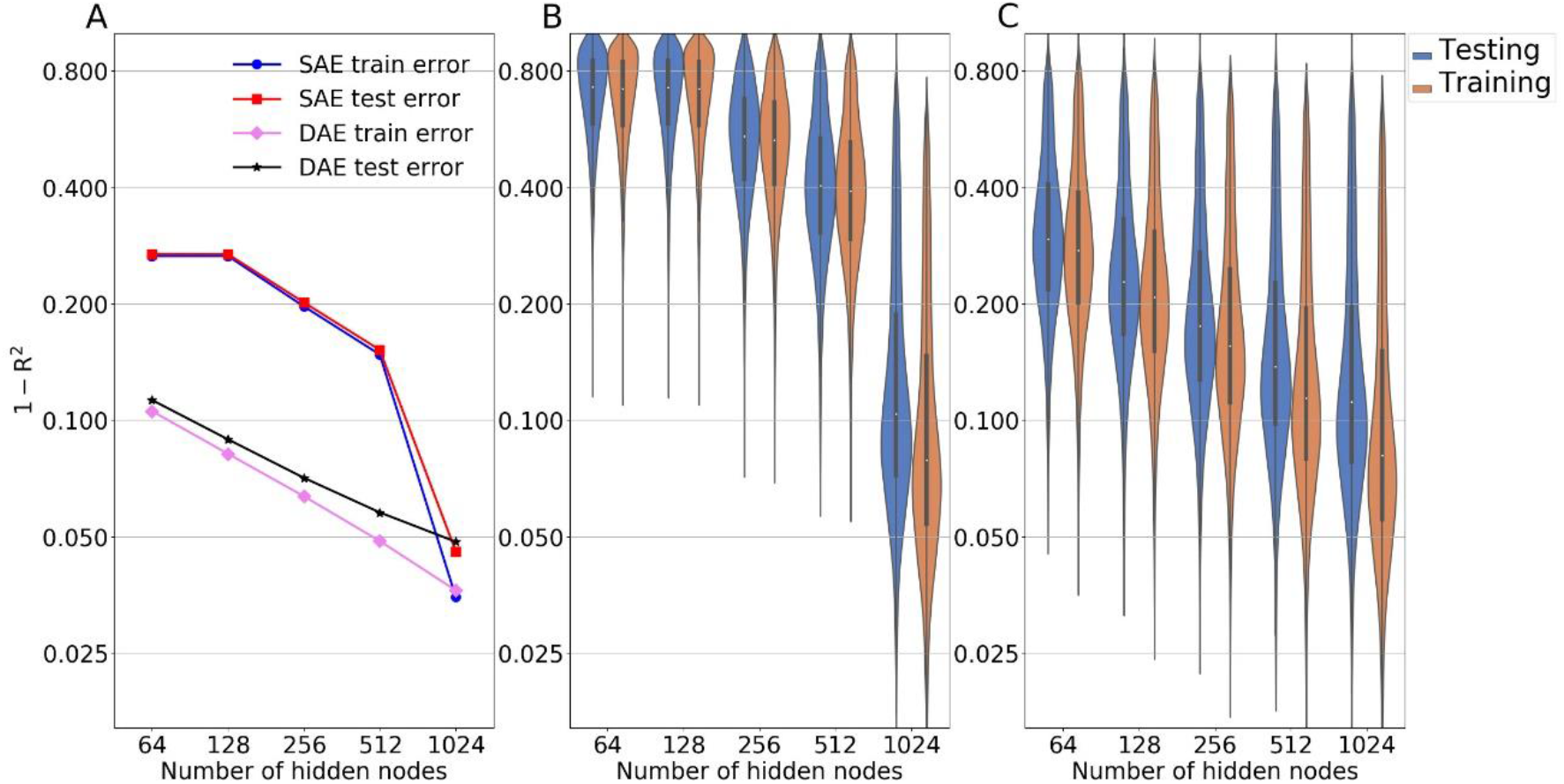
Deep auto-encoder (deepAE) outperformed shallow auto-encoder (SAE) up to 512 hidden nodes in terms of accuracy. 1-coefficient of determination (R^2^), in training and validation set using the full data set variance (A) and the gene-wise variances (B, C). Left panel shows the mean behavior of R^2^ values on the full data set. The distribution of R2 values across each gene is shown for both models, SAE (B), and 3-layer deepAE (C), increasing the number of hidden nodes in each layer from 64 to 1024.

### GWAS genes for eight different diseases were highly enriched in the third hidden layer

Our overarching aim was to assess to what extent the compressed expression representation within a deepAE could capture molecular disease-associated signatures in a data-driven manner. To this end we downloaded well characterized genetic associations for each of the diseases in our data set (DisGeNET: Piñero et al., 2017). From this data we found seven diseases in our gene expression compendium in which at least 100 genes were found in DisGeNET, which we reasoned was sufficiently powerful to perform statistical enrichment analysis. These included asthma, colon carcinoma, colorectal carcinoma, Crohn’s disease, non-small cell lung cancer, obesity and ulcerative colitis. In order to associate the genes upstream of a disease we designed a procedure which we refer to as reverse training (Methods). Briefly, using our hidden node representation and the phenotype vectors (represented as binary coded diseases) we designed a training procedure to predict the gene expression, referred to as ‘reverse’ since we explicitly used the hidden node representation. This procedure was repeated three times, (1) using only the first hidden layer, (2) using the first and second layers, (3) using all three hidden layers, and (4) as a comparison we also included the SAE.

In a result, we deciphered a gene ranking to each disease based on our functional hidden node representation. Next, we tested the relevance of this representation by overlapping the top 1000 genes of each disease with GWAS using Fisher’s exact test (Fig. 2 A). Interestingly, we found a highly significant disease association for at least one layer in all tested diseases (Fisher’s exact 10^−3^<P<10^−8^), and for five cases the strongest association was found using the full model. In all but one case (asthma) the deepAE showed a higher enrichment than the SAE. In order to validate the generality of this procedure we downloaded a new data set for MS on the same experimental platform (Brynedal et al., 2010). For this data-set we also performed a similar analysis of the control samples with other neurological disease (OND), similar to the analysis performed in (Brynedal et al., 2010). Reassuringly, we found significant enrichment for MS patients in MS GWAS (Fisher exact test P= 1.1 x 10^−5^, odds ratio (OR) = 2.3 n=29). Comparing these patients with OND patients showed lower enrichments (P= 8.6 x 10^−3^, OR = 1.7, n= 21) (Fig. 2 B) and a similar amount of top ranked differentially expressed genes between MS and OND showed no significance (P=0.49, OR=0.96, n=13). Taken together, the high enrichment of GWAS for the same disease supports our claim that our unbiased non-linear approach can indeed identify relevant upstream markers, generally with a higher accuracy than the shallower and narrower neural networks.

**Fig. (2).**
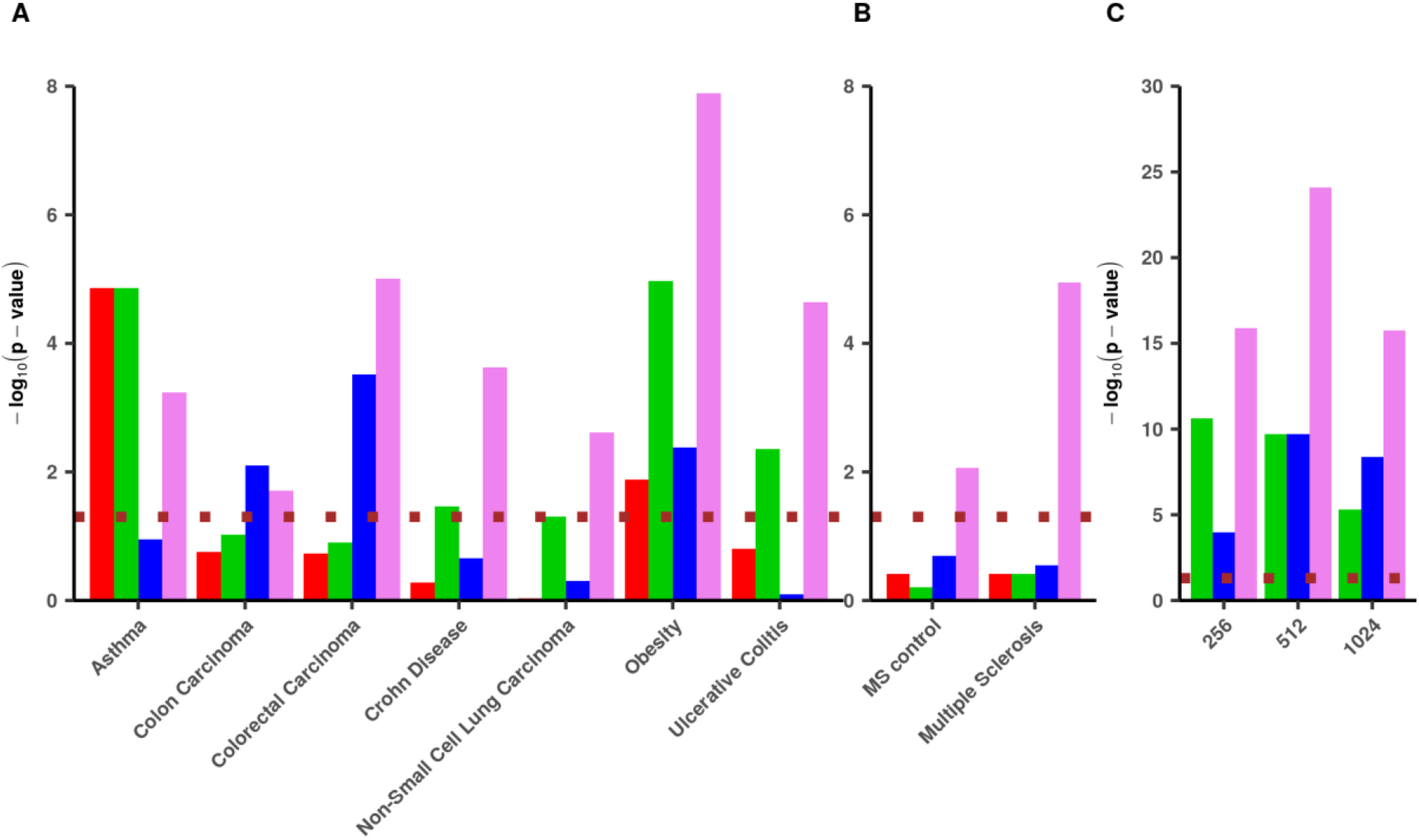
Disease association enrichment of auto-encoder (AE) derived gene sets. Enrichment score (-log10(P)) resulting from Fisher’s exact test between disease gene overlap of the predicted genes by the deep neural network derived by first (green), second (blue), and third (violet) hidden layers of the deep auto-encoder (deepAE). As a reference we also show the hidden layer of the shallow auto-encoder with 1024 nodes (SAE) that had a similar reconstruction error (red) (Panels A and B). The dotted (brown) line correspond to the p-value, cut-off 0.05 in the independent validation set in the case of control vs. MS. Panel (C) demonstrates the Fisher’s combined p-value across all the eight diseases predicted by a 3-layer deep auto-encoder.

### Samples of similar cell type and disease co-localized in the third layer of the deep auto-encoder

Next, we asked why disease genes preferentially associate with the third but not the other layers in a deep auto-encoder. However, to disentangle what is represented by each layer in a deepAE is not straightforward and has previously been the target of other studies (Chen, et al., 2016). In order to provide insight into what each layer represented in our case, we performed unsupervised clustering of the samples using the compressed representation. Since this was still a 512-dimensional analysis we further visualized the deepAE representation using the first two linear principal components (PCs) of the compressed space. This representation is henceforth referred to as the deepAE-PCA. Previously it has been shown that classical PCA on the full ~20,000-dimensional gene space can discriminate cell types and diseases very well, which we therefore used as a reference in our analysis (Torrente et al., 2016). To analyze whether samples close in these spaces were biologically more similar than two random samples, we computed the Silhouette index (SI) for phenotypically defined groups, governed by their cell type and disease status respectively (Fig. 3). Note that SI=1 reflects a perfect phenotypic grouping and SI=-1 indicates completely mixed samples. Next, the samples were grouped based on the different cell types in the data (n=56) and tested to determine whether the deepAE-PCA or PCA had the highest SI based on each of their respective, different hidden layers (see Methods). We filtered the compressed coordinates of normal cell types and found significantly more cell types having a higher SI for the third hidden deepAE layer than was the case in the PCA-based approach (n=45 out of 56, odds ratio = 4.09, binomial test P= 5.4×10^−6^). Interestingly, smaller enrichments were also found for the first (n=37, OR= 1.95, P=0.022) and second (n=37, OR=1.95, P=0.022) layers. Next, we repeated this analysis for the 128 diseases we had in our data, and we found all the cases showed strong associations: first layer (n=97, OR=3.13, P= 4.2×10^−9^), second layer (n=98, OR=3.27, P= 1.3×10^−9^) and third layer (n=98, OR=3.27, P=1.3×10^−9^). These observations suggested that samples originating from similar conditions and phenotypes were automatically grouped according to the hidden layers, most significantly for the third.

**Fig. (3).**
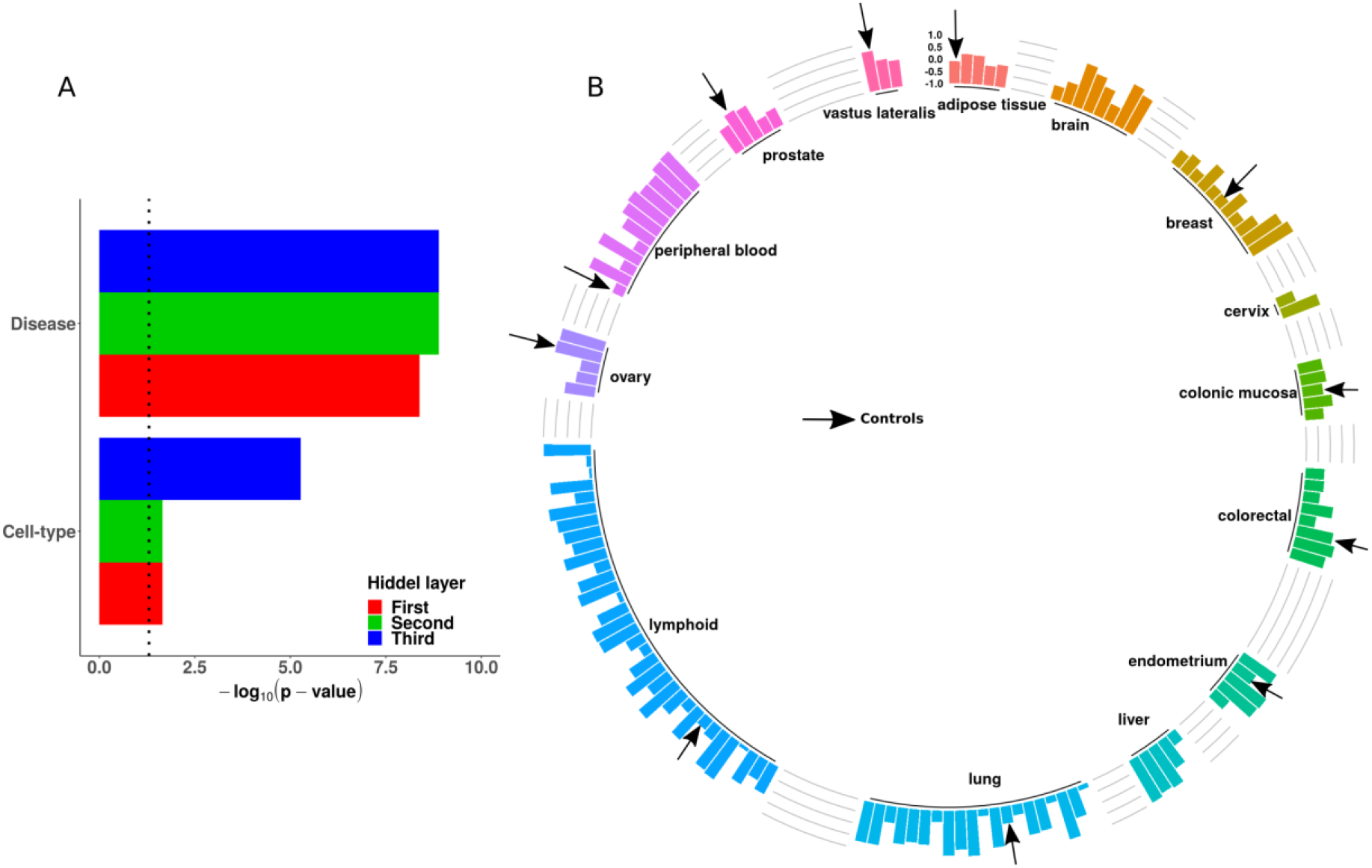
Deep auto-encoder (deepAE) representation clustering samples into cell types and diseases. (A) Significance score (-log10(p)) for first (red), second (green), and third (blue) deepAE layers are more coherent (measured by a high Silhouette index (SI)) with respect to cell types (lower) and diseases (upper) than the standard principal component (PC) analysis-based approach. (B) SI defined by the two PCs for diseases and control samples on compressed signals at the third hidden of deepAE with each of the three hidden layers having 512 nodes. Control samples are marked by arrows.

### Genes associated with the same hidden node in the first layer were co-localized in a protein-protein interactome

In order to further interpret the different layers and uncover their role in defining disease modules, we proceeded to analyze the relationship between the signature genes of each hidden node. Since cellular function is classically described in terms of biological pathways, or lately has also been abstracted to densely interconnected sub-regions in the interactome (so-called network *communities*) we analyzed the parameters in the deepAE and their connection to the global pattern of expressed genes (Amorim et al. 2018; Lin et al., 2017). There are different ways one could potentially interpret parameters in a deepAE. To this end we, created a procedure to associate genes with hidden nodes which we refer to as *light-up*. Briefly, a light-up input vector was defined for each hidden node by activating it to the maximum value of the activation function, clamping all other nodes at the same layer de-activated by zero values. Then we forward-propagated this input vector through all layers to the output pattern response on the gene expression space (Methods). This resulted in a ranked list of genes for each hidden node, identifying which genes were most influenced by the activation of that node. We repeated this procedure for all hidden nodes and layers. In order to test if these lists corresponded to functional units, we analyzed their localization within the protein-protein interaction network STRING (Franceschini et al., 2013, ver 9.1). We hypothesized that genes co-influenced by a hidden node could represent protein patterns involved in the same function. Also, the STRING database captures proteins associated with the same biological function and which are known to be within the same neighborhood of their physical interactome. By first ranking the most influenced genes we systematically analyzed the cut-offs thereby showing whether a gene was considered as associated with the node by powers of two from 100 to 10,000. Next, we calculated the average shortest path distance between these genes within the STRING network, using the harmonic mean distance to include also disconnected genes. This analysis revealed that the top-nodes in the ranked lists of the first hidden layer, had a high betweenness centrality (Fig. 4A) while exhibiting a low average graph distance between each other (Fig. 3B-D). Thus, highly co-localized genes were the most central part of the PPI. Both findings were tested using several different cut-offs (Fig. 4A-D) and the effect was most evident for the first layer, appearing to a weaker extent for the second layer and fully vanishing at the third layer.

**Fig. (4).**
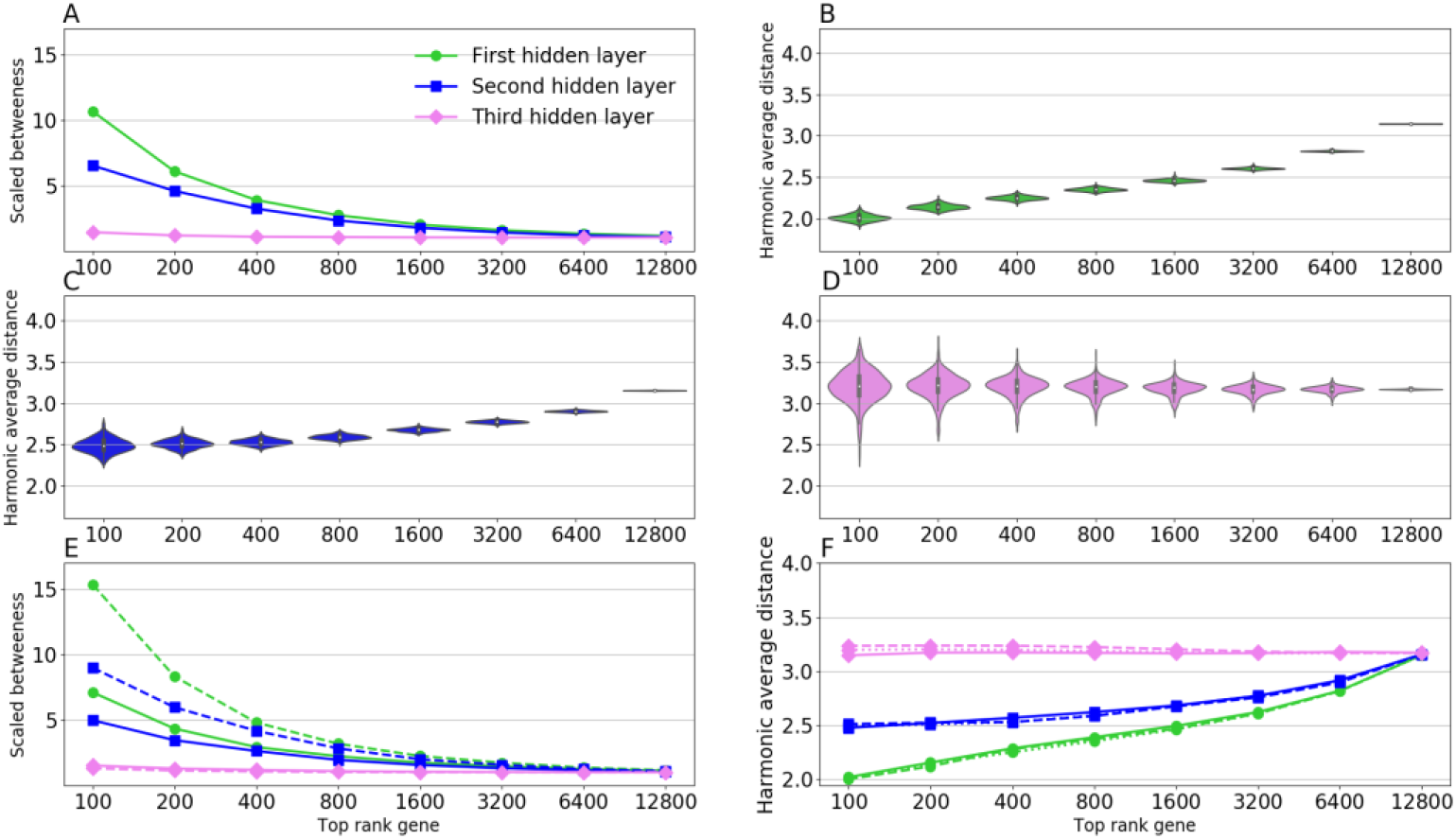
Panel (A) demonstrate the betweenness centrality behavior of the top ranked genes on the basis of the first (green), second (blue) and third (violet) hidden layers of the deep auto-encoder. Panels (B), (C) and (D) show the distribution of harmonic average distances of the top rank genes based on each hidden node of the first, second and third hidden layers of the deep auto-encoder respectively. Also, these results are robust across 256 and 1024 hidden nodes of the deep auto-encoder (Panels (E) and (F)).

### Validation of the approach using RNA-seq transcriptomic data

In order to assess the generality and to increase the domain of applicability of the auto-encoder approach to interpret emerging large RNA-seq data sets, we identified a large publicly available body of RNA-seq material (Lachmann et al. 2018). This data was divided into 50,000 training samples and 9,532 validation samples for 18999 genes, and was used to train a deep AE with similar hyperparameters as for the microarrays, i.e., using a three-layered AE with 512 hidden nodes in each layer. Unfortunately, this data did not contain sufficient complex disease samples and we therefore searched for additional RNA-seq data sets for our previously tested complex diseases, namely asthma (GSE75011), Crohn’s disease, ulcerative colitis GSE112057, obesity (GSE65540) and multiple sclerosis (James et al., 2018). Similar to the microarray AE we found a highly consistent significant overlap between GWAS and the associated disease genes derived from the third layer for each of the diseases (Fisher combined P<10^−12^), and to a lesser extent in the other two layers, see Fig. 5. Next, we tested whether the hidden nodes corresponded to close sets of interconnected protein-protein interactions by repeating the light-up procedure. Interestingly, we found that the top ranked genes in the first, and to a lesser extent also in the second hidden layer, had low average betweenness centrality and had low average distance. Strikingly, this association was even stronger than in the analysis using the AE of the microarrays. In summary, our replication of our findings that the relationship between disease gene and the protein interaction confirms our findings of deep AEs as an unbiased estimator of functional disease associations (Fig. 6).

**Fig. (5).**
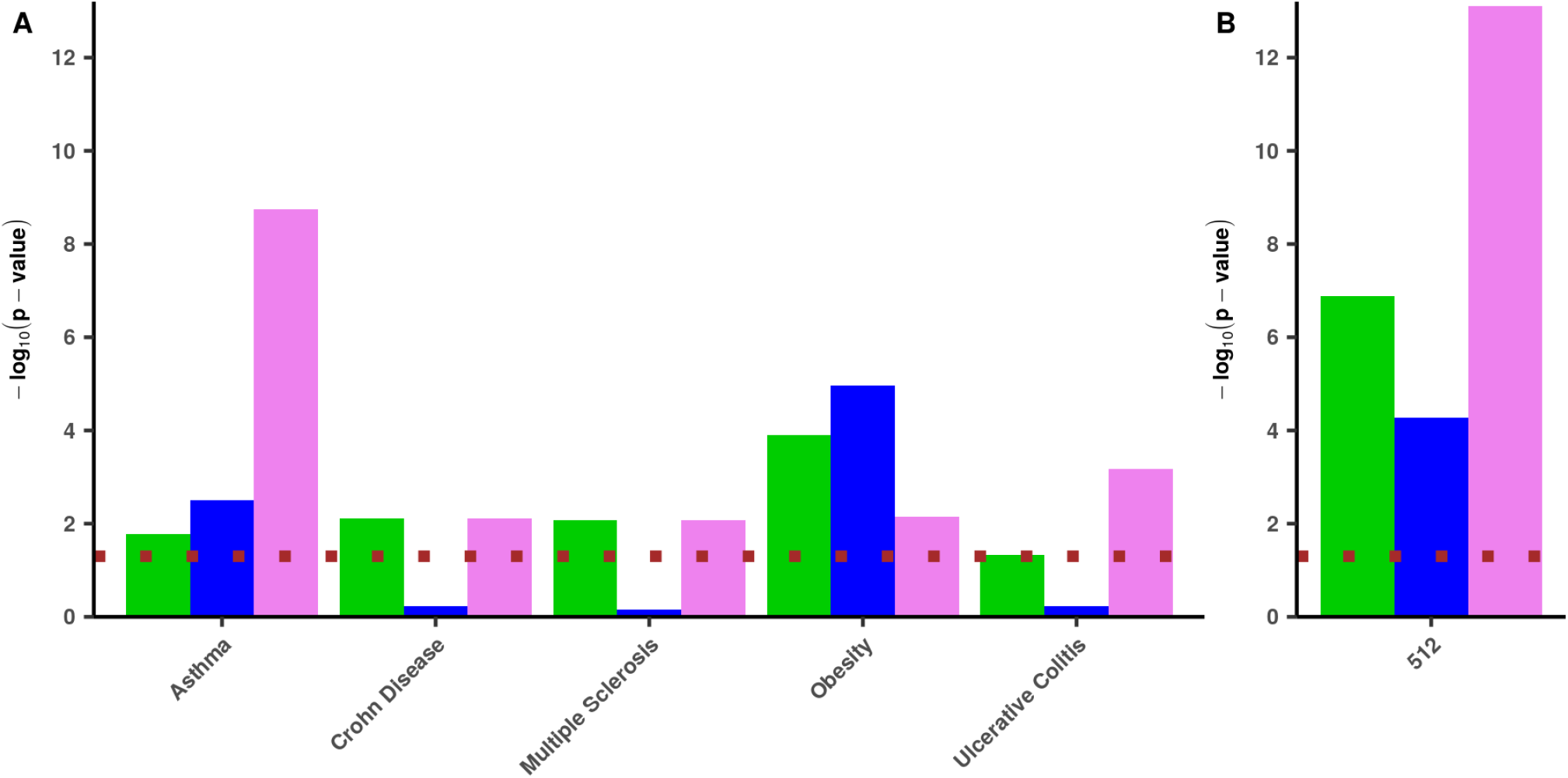
Validation of disease association enrichment results of deep auto-encoder (deepAE) derived gene sets on RNA-seq data. Enrichment score (-log10(P)) resulting from Fisher’s exact test between disease gene overlap of the predicted genes by the deep neural network derived by the first (green), second (blue), and third (violet) hidden layers, of the deepAE in the Panel (A). Panel (B) demonstrates the Fisher’s combined p-value across all the five complex diseases predicted by the 3-layer deep auto-encoder. The dotted (brown) line corresponds to the p-value, cut-off 0.05.

**Fig. (6):**
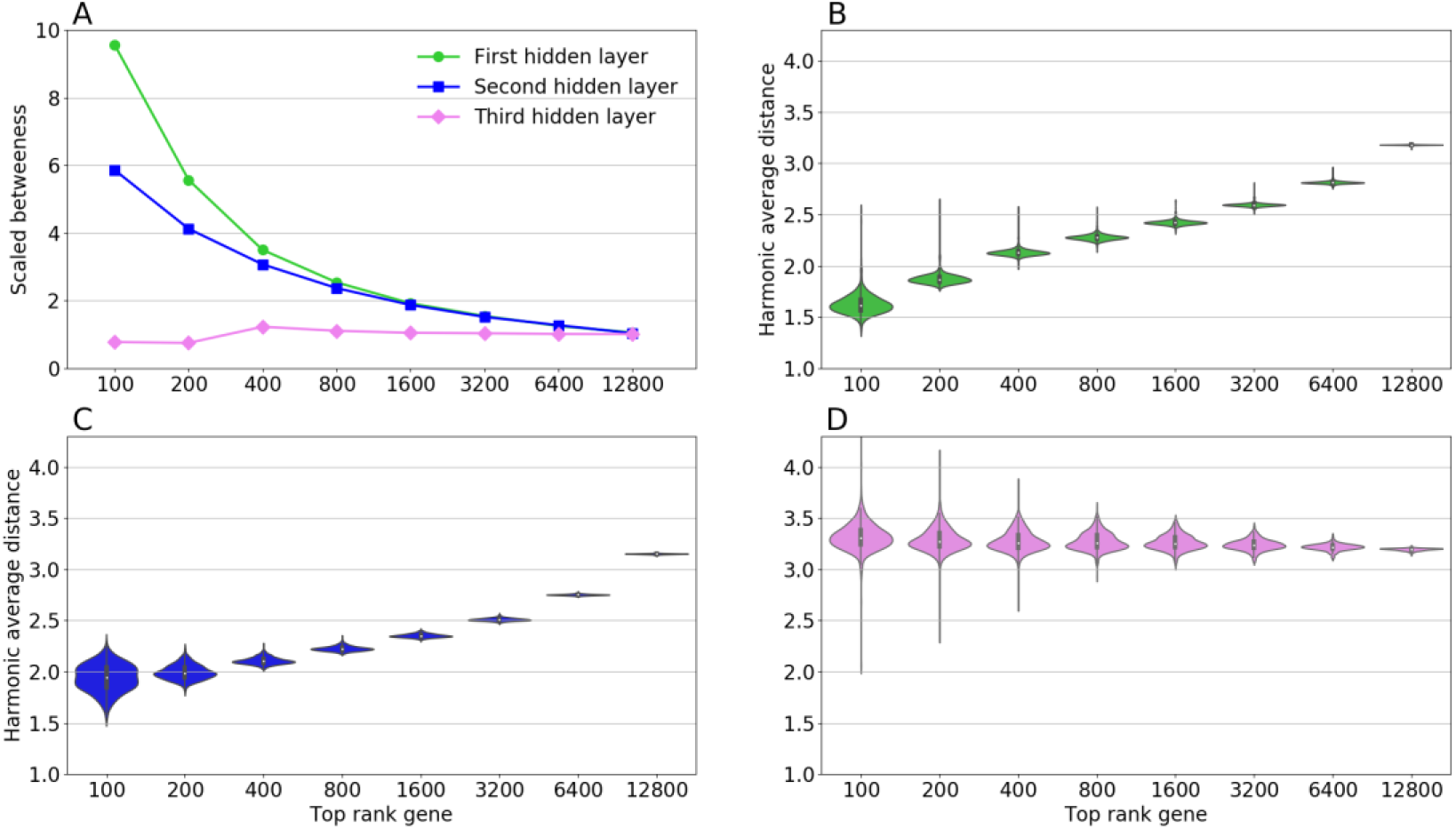
Panel (A) demonstrates the betweenness centrality behavior of the top ranked genes on the basis of the first (blue), second (red) and third (pink) hidden layers of the deep auto-encoder trained on the RNA-seq data. Panels (B), (C) and (D) show the distribution of harmonic average distances of the top rank genes based on each hidden node of the first, second and third hidden layers of the deep auto-encoder respectively.

**Fig. 7.**
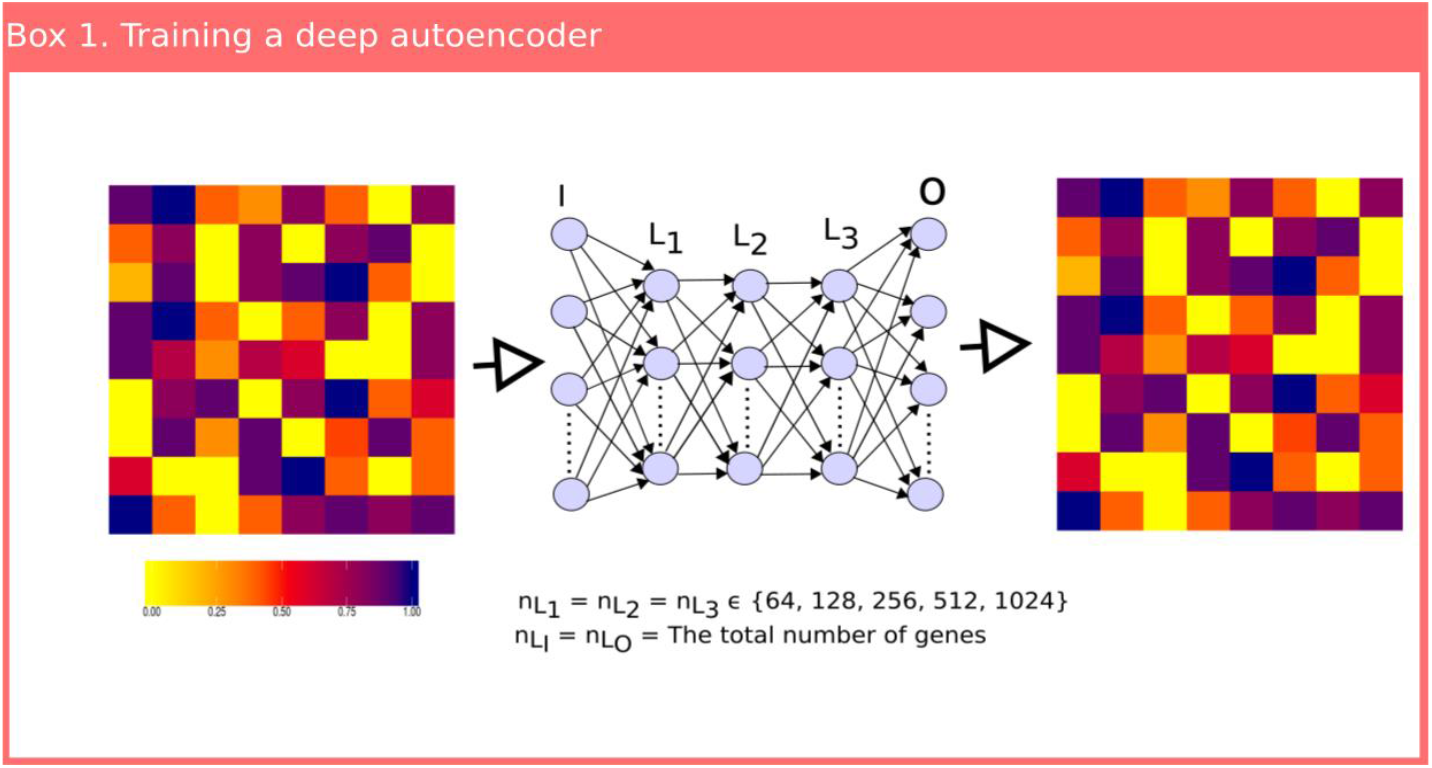
Schematic diagram of training an auto-encoder from normalized transcriptomic data.

**Fig. 8.**
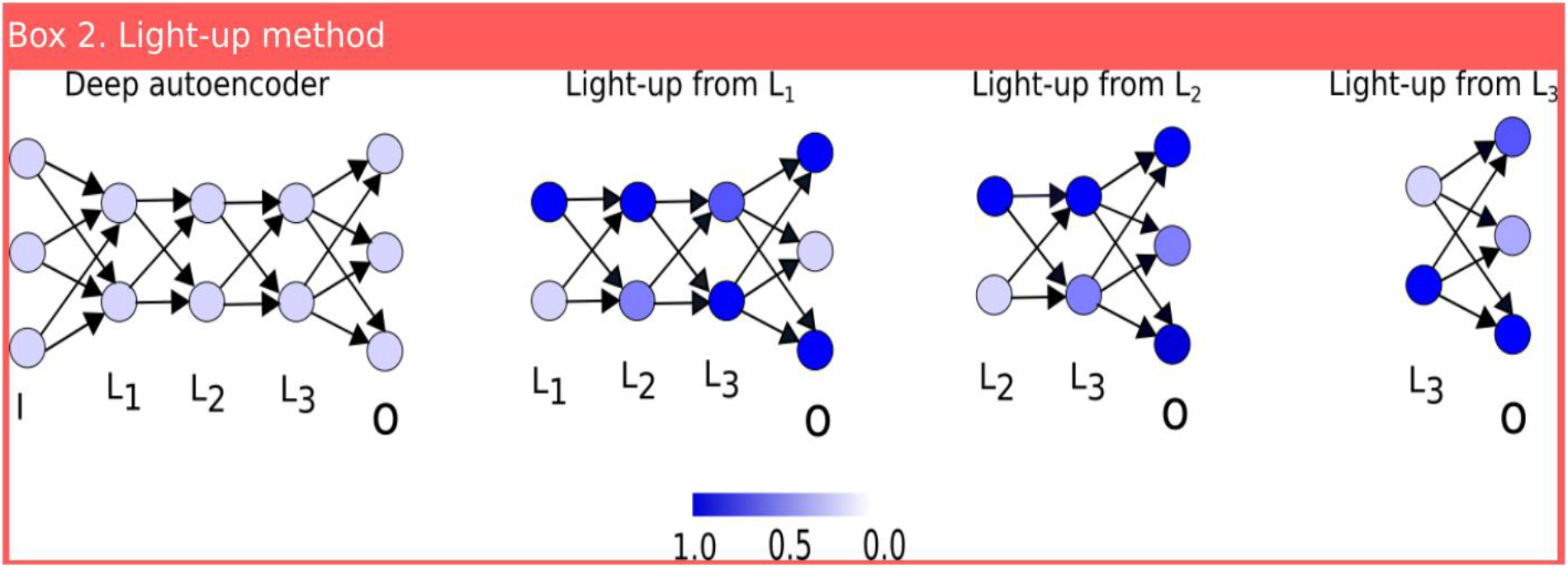
Schematic diagram for interpreting auto-encoder in terms of the PPI. Description of the light-up from the first, second and third hidden layers by the trained deep auto-encoder.

In the last few decades, various interactome connectivity-based approaches have been proposed for defining the disease module; however, the consistency of such approaches across the various diseases is limited due to incompleteness and the biased nature of the interactome (Menche et al., 2015). On the other side, the increase in the size of available transcriptomic data could offer an opportunity to overcome this method-biased problem by finding the functionally-related molecular components. The method represents a step toward overcoming the drawback of network method to biased underlying networks.

## Discussion

In summary, our study aimed at using deep neural networks for identification of unbiased new functional representation that can explain complex diseases without the reliance of the protein-protein interaction (PPI) network which is known to be incomplete (Menche et al., 2015) and strongly affected by the study bias of some early discovered cancer genes. We showed for the first time the applicability of deep learning that could prevail over the findings derived from network medicine for understanding complex diseases. In order to find the similar inferences between structural features of the PPI and the estimated parameters of neural networks, we began a systematic demonstration of the *light-up* concept (Mordvintsev et al., 2015) motivated by the need to prioritize genes based on their contributions in the compressed space of the deepAE. Furthermore, we showed that the top genes prioritized by each node in the first and middle layers are localized and belong to the core part of the PPI. Moreover, the third layer nodes possess long-range variability in showing the localization to de-localization of their top genes compared to the random genes. This kind of gradient in terms of interpretability with respect to localization within the PPI network suggests that each layer indeed encodes different types of biological information. These results also suggest that the transformed signals in the compressed space first decode the modular features of the underlying interactome which then vanishes smoothly layer-by-layer as a deeper representation is encoded. Concurrently, with such a decreasing protein-defined modular gradient, an increase in disease-relevant genes and modules thereof is progressively discovered in the deepest layers of the auto-encoder. Next, we presented a novel method that follows a neural network to determine a disease-specific feature vector in the compressed space of the deepAE. The disease-specific feature vector of the compressed space transformed to a gene space that defined the disease module. For this purpose, we found deep auto-encoders using 512 state variables of ~20,000 genes at a 95% R^2^ for microarrays and 80-85% for RNA-seq. This represented a two-fold compression compared to the variables in the LINCS project and to our tested shallow. The high degree of compression for the deep auto-encoder with fewer state variables suggests that this representation is indeed preferable compared to shallow representations. One reason for the need of deepness is that such AEs are theoretically capable of capturing more complex relations between genes, such as the XOR relations (Goodfellow et al. 2016, Hong et al. 2013, Hunziker et al. 2010), which shallow AEs cannot.

Our findings suggest the usefulness of deep learning analysis to decompose different hierarchy levels hidden with the relations between genes. For example, the first layer encodes the modular features belonging to the central part of the interactome. These features are synonymously selected by the interactome-based approaches to find the components that have control over the entire system (Wuchty, 2014). In contrast, these features are not necessarily transferable by cell type-specific transcriptomic signals. More interestingly, the third layer efficiently encodes cell type-specific functional features; therefore, it might be reason to increase the likelihood of mapping the disease-specific functional genes by disease-related cell type signals in the light-up. Also, the presented approach can play a crucial role in utilizing the resolution level of the single cell transcriptomic signals in prioritizing genes that are enriched with the upstream dysregulated genes and their relationship with causal genetic variants (Calabrese et al., 2017). Another important application of our approach can indeed provide new insights in the multi-scale organization about disease-disease, disease pathways disease-gene associations (Gaudelet et al., 2019).

Using transfer learning our AEs could help stratify disease groups of limited samples as the number of parameters could decrease by about ~40-fold (from ~20,000 to 512), which decrease the analysis complexity. Therefore, transfer could be applied by other clinically interested researchers staring from our derived representation, which could lead to increased power for building classification systems. Lastly, we showed the generality of our approach by confirming our result in RNA-seq data. We think the approach is applicable to other omics and using our derived single omics representations together with others they open a door to multi-omics neural networks using transfer learning, similar to what is nowadays routinely done within the field of image recognition.

## Methods

### Auto-encoder construction

#### (1) Data preparation and normalization

The micro-array data is log transformed normalized values. Similarly, we normalized RNA-seq data by the upper quantile method using the function uqua of the R package NOISeq and log transformed the normalized gene expression values by log_2_(1+normalized expression value) (Tarazona et al., 2011). Also, we discarded the noisy log transformed expression values those are less than the 3.0. Next, we re-normalized the *i^th^* gene mRNA expression level in the *k^th^* sample 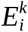 across the samples to be in the range between zero and unity, such that

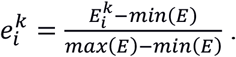

#### (2) Parameter optimization

The normalized expression matrix 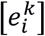 is input and output signals for training the auto-encoder with sigmoid activation function. We have chosen a dense layer so that the optimizer starts with an initial point that has unbiased dependency among the data features. We used optimizer ADAM, with learning rate = 0.0001, *β*_1_ =0.9, *β*_2_=0.999, *ϵ* =1e-8 and decay =1e-6, to train the model which we have observed as an optimal choice in predicting the high level of accuracy in the both training and validation data sets (Kingma, D. P., and Jimmy Ba (2014). The batch size was 256 for the training. In order to systematically investigate the impact of number of hidden nodes on the prediction accuracy, we fixed the number of hidden nodes in all the three hidden layers of the deep auto-encoder (deepAE), termed as a three-layer model. In our case, the three-layer model with 512 hidden nodes was more suitable for capturing the biological features. This model has fewer reconstruction errors in comparison with similar hidden node of the one-layer model (SAE). We implemented our methods using the tensorflow backend (https://www.tensorflow.org) and Keras (https://github.com/keras-team/keras) neural network Python-library.

### Interpreting the trained auto-encoder with PPI

The preserved biology in the compressed space is confined in each hidden layer. Therefore, our objective was to understand the meaning of all the nodes in each hidden layer. For this objective, we computed activations at the output layer for each node of a hidden layer. We recursively forward-propagated the maximum activation value of each node, while keeping other nodes neutral by zero input, on the remaining portion. Finally, we prioritized the genes on the basis of last layer activations. For simplicity, we mathematically formulated these steps as follows. Suppose k^th^ layer of an L layer AE, has N^k^ nodes. Here, N^1^ and N^L^ are the same as the number of genes in the profile expression matrix. Also, the number of nodes in each hidden layer is h, i.e. N^k^ = h for 2≤k≤L-1. The following equation recursively defines the activations, x^k^, of the k^th^ layer from the activations, x^k-1^, at (k-1)^th^ layer with the initial activation vector x^p^ (it consists of the maximum activation value at the corresponding position of the hidden node and the rest of the elements are zero) corresponding to the node in the p^th^ hidden layer,

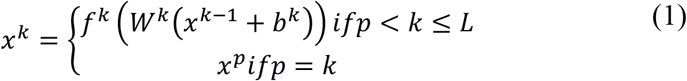

Where *f^k^*, *b^k^* and *W^k^* are associated with the k^th^ layer activation function, bias term and weight matrix respectively. Note that the first input layer does not have an activation function, bias term and weight matrix, so 2 ≤ k ≤ L. The equation (1) defines the activations at the output layer, *x^L^*, with dimension of gene size. We prioritized the genes based on the vector, *x^L^*, to show the associations with the PPI module.

### Predicting disease genes

We derived a new approach for predicting a disease gene that is explained in the following four steps (Fig. 9):

**Fig. 9.**
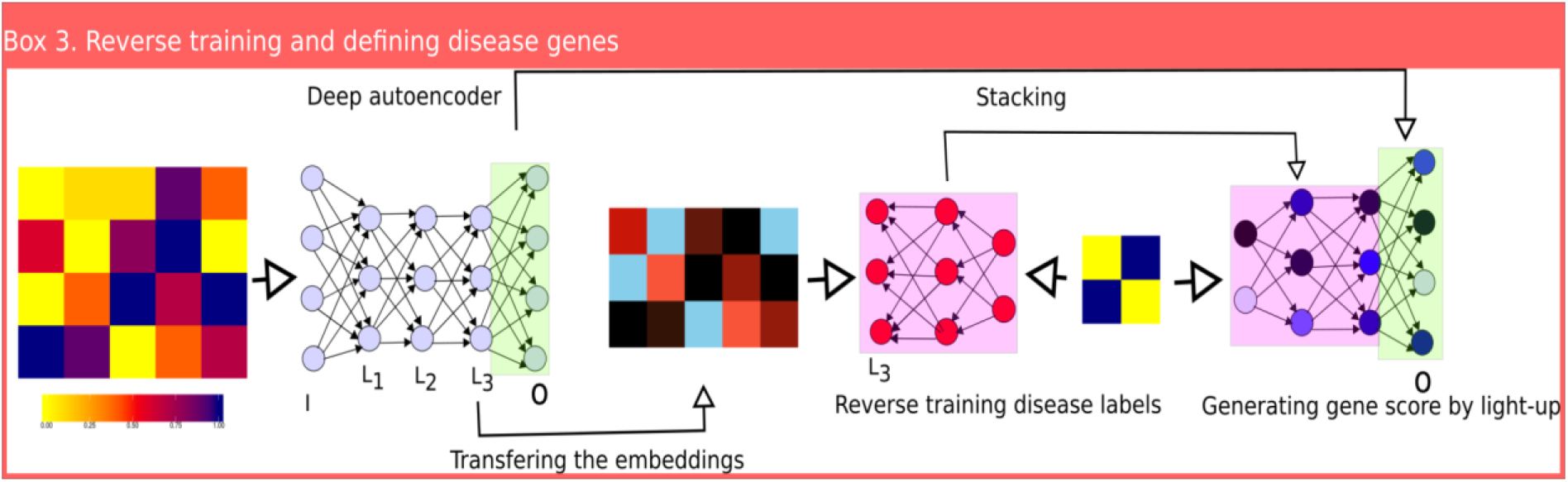
Schematic diagram illustrating making disease associations using the trained three-layer deep auto-encoder and the deep neural network-based method. Most of the left colored matrix corresponds to the normalized gene expression profile where rows and columns are associated with genes and samples, cells and corresponding phenotypes, respectively. The next colored matrix corresponds to the compressed representation of the expression profile at the third layer by the deep auto-encoder.

**(1) Compressing the expression profile** at hidden layers using trained deepAE.

**(2) Training a supervised neural network:** We trained a one-hidden-layer supervised neural network, having the same number of nodes in the second and third layers, with sigmoid and linear activation function respectively. The input matrix 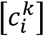 is followed by 1 ≤ *k* ≤ *S* and 1 ≤ *i* ≤ *P* with dimension *S*x*P*, where *S* and *P* are the total number of samples and their associated phenotypes respectively. The matrix 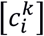 is defined by another identity matrix 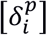 of the Kronecker tensor as follows, 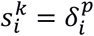 if the *k*^th^ sample is associated with the *p*^th^ phenotype. The output matrix 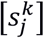 is a profile matrix of compressed signals at a hidden layer of dimension *S*x*h*, while *h* is the number of nodes in the hidden layer.

**(3) Stacking the supervised neural network** with the left part of deepAE, in the feed forward direction, from the layer at which the supervised neural network is trained. We scaled the mean and the variance of the weight matrices and biases in the consecutive layers where the both networks are stacked.

**(4) Finding the disease scores from the expression:** The scores s^p^, for prioritizing the genes related to the p^th^ phenotype are computed by the parameters of a stacked neural network using:

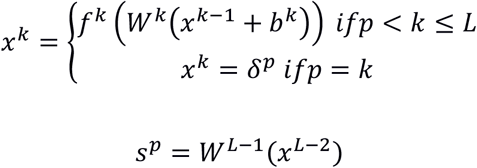

Where *δ^p^* is a Kronecker tensor for the p^th^ phenotype.

### Validation of predicted genes

We downloaded the curated disease SNPs from the DisGeNET database and human genome reference consortium assembly, build version 37 (GRch37, hg19) from the UCSC database (https://genome.ucsc.edu/). We computed the closest gene to each disease associated SNP, using Bedtools under the default option. In this way, we defined disease-associated gene sets for validating the neural network-based predicted genes. The performance of the predicted genes was demonstrated in terms of Fisher p-value using a hypergeometric test.

